# Architecture of a multi-cellular polygenic network governing immune homeostasis

**DOI:** 10.1101/256073

**Authors:** Tania Dubovik, Elina Starosvetsky, Benjamin LeRoy, Rachelly Normand, Yasmin Admon, Ayelet Alpert, Yishai Ofran, Max G’Sell, Shai S. Shen-Orr

## Abstract

Complex physiological functionality is often the outcome of multiple interacting cell-types, yet mechanistically how a large number of trait-associated genes yield a single multi-cellular network governing the phenotype has not been well defined. Individuals’ immune-cellular profiles at homeostasis show high heritability and inter-individual variation with functional and clinical implications. We profiled immune cellular variation by mass-cytometry in 55 genetically diverse mouse strains. We identify 788 genes associated with cellular homeostasis, supporting a polygenic model where 52% of genes correspond to core homeostatic functions whose genetic variants suffice to predict phenotype. Trait genes form a multi-cellular network architecture showing increased functional complexity over evolutionary timescales for shared regulation to all cells, specialized cell-specific programs, and between-cell synchronization. Contrasting to human studies suggests the regulatory network expands with environmental exposure history. Our findings shed light on the origin of immune-cellular variation and regulatory architectures that may generalize to other environmentally sensitive systems.

## Introduction

Complex physiological functionality is often the outcome of multiple interacting cell-types. Thus, understanding of genotype-phenotype relation requires cell-specific resolution of regulatory networks as well as an understanding of the interaction between cell-types. QTL studies have yielded large advances in mapping the genotype-phenotype connections, with eQTL studies likely being the most common means of assembling regulatory networks linked to genotype. However, these studies often rely on whole tissue (‘The Genotype-Tissue Expression (GTEx) pilot analysis: Multitissue gene regulation in humans’, 2015) or sorted cell profiling where the multi-cellularity context is absent (Heng and Painter, 2008; Dimas *et al.*, 2009; Link *et al.*, 2018; Schwartzentruber *et al.*, 2018). Most recently, profiling single cell gene expression and genotype of multiple cell-types has become a viable option (van der Wijst *et al*., 2018), yet due to prohibitive costs and high level of noise, generating networks and associating their variation to genotypic variants remains difficult. Moreover, as mRNA is the studied trait, physiological context remains missing.

Cellular immune profiles are a complex physiological trait reflecting cell-type abundance for each cell in tissue and showing a large amount of variation between individuals with important diagnostic value in health and disease (Gaudillière *et al.*, 2014; Tsang *et al.*, 2014; Brodin and Davis, 2017), suggesting this complex system maintains a “personalized immune homeostasis”. Cellular immune profiles are highly heritable, which may be due either to genetics, shared environmental influences or a combination of both (Lu *et al.*, 2016). Repeated evidences from young mono- and di-zygotic twins studies suggest a high level of genetic determinism (Evans, Frazer and Martin, 1999; Hall *et al.*, 2000; Pedersen, 2000).

From a functional perspective, evidence from basic studies tracking labeled cells have shown that when one changes the rate of a cell subsets’ proliferation or death, the total number and relative frequencies of cells would be altered (Mohri *et al.*, 1998; Asquith *et al.*, 2002; Busch *et al.*, 2015). Extrapolated from these experiments, and fitting with mathematical modeling, a cell abundance should be a function of its turnover. Namely, a balance of processes involved in the introduction of new cells (e.g. proliferation, or influx of migrating cells) and processes involving in cell clearance (e.g. death, differentiation to another cell-type, or an outflux of migrating cells). Yet, several large-scale association studies that were conducted in humans found only a few genes associated with blood cell subset frequencies and counts (Orrù *et al.*, 2013; Roederer *et al.*, 2015; Patin *et al.*, 2018), making it difficult to discern a genetic architecture for those associations identified. This may be partially explained by data from mono-zygotic twins showing that the effects of the environment predominantly increase with age (Brodin *et al*., 2015) and therefore detection of genetic associations may become more difficult.

Particularly of relevance, recently it has been proposed that though all genetic variants play a role in complex trait determination, a modest subset of ‘core genes’ exist which directly affect the trait and are related to its biology (Boyle, Li and Pritchard, 2017). This core genes hypothesis has been contested, particularly with respect to the size of gene group, which was hypothesized to extend to include all genetic variants in the genome (Wray *et al.*, 2018). In case of immune cell abundance, the trait is well defined, and the genes associated with its core biology, in theory, should be well defined as well.

We performed an association study of bone marrow immune cells in Collaborative Cross (CC), a panel of highly genetically diverse inbred mouse strains for which a tailored SNP chip is available along side the full genome sequence of all founder strains (Consortium, 2004). The CC exhibit large phenotypic variation with respect to immune homeostatic composition (Kelada *et al.*, 2012; Graham *et al.*, 2017), thereby allowing studies with statistical power, and reproducibility in a setting with minimal and equal environmental influences on immune cellular homeostasis. We show that baseline variability in immune cell profiles is determined by a large number of genetic variants. We use these variants to infer a multicellular regulatory network whose associated genes form protein-protein interactions within cells as well as inter-cellular interactions determining immune cellular homeostasis. We characterize the architecture of this mouse-derived network through an evolutionary lens as well as how it is reflected in environmentally exposed humans, allowing inference of polygenic network dynamics.

## Results

### Highly variable immune cellular profiles in genetically diverse mice are polygenic complex traits

We focused on the bone marrow, given it covers major developmental milestones of hematopoiesis. The CC mouse panel was designed to achieve individual genetic homozygosity from eight inbred founder strains while maintaining heterogeneity between strains (founder strains consisting of five classical inbred and three wild-derived, with roughly 75% of variation stemming from the latter). Fitting the need for profiling this complex cellular environment at high dimension, we profiled 30 naive CC mouse strains in duplicate and 8 founder strains in triplicate (**Table S1**) using a large panel of 20 CyTOF phenotypic markers, reflective of hematopoietic populations (**Table S2**). We noted that for several strains, some markers were unobservable (**Table S3)**. Considering the remaining samples, which had a full set of phenotypic markers (15 CC strains and 3 founder strains), we clustered their single cell data using a high-dimensional clustering algorithm (**Figure 1A**, **Table S4**, see Methods) (Bruggner *et al.*, 2014). This resulted in immune profiles of CC mice being clustered by replicate strains, followed by a close similarity to the whole genome genetic kinship distance, suggesting that phenotypic similarity may be driven by genetic similarity (**Figure S1A**). To assess the variation between individual mice’s homeostatic composition (a vector of multiple cell subset frequencies) we performed a Principal Component Analysis (PCA) of bone marrow cell subset composition (**Figure 1**B). We noted that the first two principle components axes explained 48.3% of total variance, with the first component capturing variation in lymphoid lineage B and T cell subsets and the second component capturing variation in myeloid cell subsets and NK cells. To quantify the extent of variation in homeostatic composition, we measured the pairwise distance between individual mice profiles within replicate mice, founder and CC strains. CC strains created a continuous scale of phenotypes likely introduced by synergistic genetic interactions, whose dynamic range was largest and mean distance significantly greater than that observed between replicate strains (**Figure 1**C, p<0.001 by *Student’s t*-*test*). Interestingly, we noted no significant differences of homeostatic composition distances between bone marrow samples profiled from ten healthy adults and CC strains, supporting the observation that these strains phenotypic variation is close to that observed in humans (**Figure 1**C, **Figure S1**B for human gating scheme) (Rogala *et al.*, 2014; Elbahesh and Schughart, 2016). Taken together, we demonstrate that immune cell subset variation is high, yet non-random, in genetically diverse mice raised in a shared clean environment.

**Figure 1.**
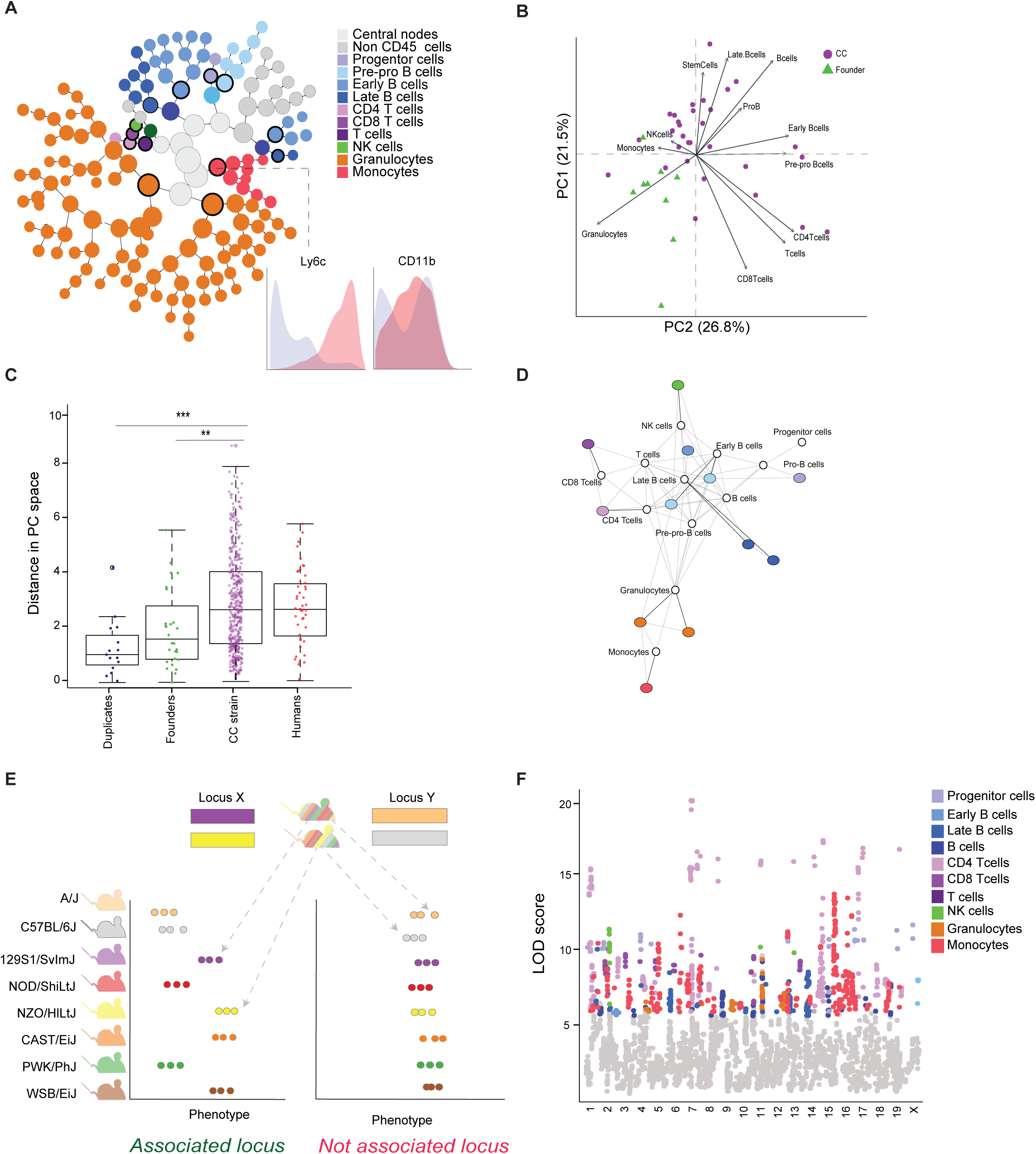
Immune cellular profiles are highly variable across genetically diverse CC mice strains and their regulation is polygenic. **(A)** Tree structure of Citrus high dimension clustering of single cell bone marrow hematopoietic cells from Collaborative Cross mice. Cell subsets are color-labeled according to phenotypic markers distinctive for major bone marrow populations. Circled nodes were used for further downstream analysis. Inset shows example of Ly6c and CD11b single cell marker expression distributions (red and blue, cluster versus background respectively), contributing to cluster characterization of monocytes. **(B)** PCA of immune cellular profiles obtained from manual gates of CC (purple) and founder (green) strains. Only strains with complete set of markers were included (**Table S3**). **(C)** CC strains form a continuous phenotypic variation in immune homeostatic profiles whose dynamic range exceeds that of founder strains and is similar to that observed in humans. Boxplots showing pairwise analysis of distances in PC space (the first and second dimensions were used) within groups of duplicates, founder or CC strains as well as ten humans healthy bone marrow. Student’s t-test p-value are: ^∗∗^p<0.01, ^∗∗∗^p<0.001. Pairwise distances between replicate strains appear in the Duplicate group. **(D)** Scaffold mapping between manually gated populations and those obtained from Citrus clustering analysis overcomes heterogeneity in marker expression. **(E)** Illustration of QTL mapping in CC mice: CC strains genome is reconstructed at each locus to one of eight founder strains are assayed for a quantitative trait such as an immune cell subset frequency. For each locus, and each CC strain, mice are split based on which founder strain contributed the locus (e.g. ‘Locus X’ for the left mouse is contributed by 129S1/SvImJ and for the second mouse by NZO/HlLtJ), we then test phenotypic association across all founders. **(F)** Manhattan plot for all associated loci, the threshold of association is set according to FDR of each population. Colors denote cell subset gene was mapped to.

Next, we sought to link phenotypic variation to genotype. To increase our statistical power we deployed several strategies: First, we leveraged the Scaffold algorithm which avoids bias in cluster identity by mapping high dimensional clusters to manually gated populations using the same set of markers, thereby allowing to increase the mouse samples size (Spitzer *et al*., 2015) (**Figure 1**D, **Figure S1**C for gating scheme, see Methods). Second, we focused on loci harboring genes expressed in immune cells and for which coding region functional mutations were predicted (**Table S5**). This filtering procedure yielded 6,961 genes covering a broad set of functions, spanning from broad cellular to immune specific functionalities. Next, we mapped per mouse loci, the likelihood of it stemming from each founder strain, using haplotype reconstruction (see Methods) and computed the logs odds ratio (LOD) score of each locus with each of the 11 manually gated cell subset populations (**Figure 1**E). This procedure yielded a total of 1,617 loci, corresponding to 1,579 genes which matched our filtering criteria, namely below a false discovery rate (FDR) threshold of 5% and robust to a leave-one-out procedure (**Figure 1**F, **Figure S1**D, **Table S6**, see Methods). Thus, a large number of genes are associated with this multi-cellular complex trait.

### Genetic associations differ by cell-type and include inter-cellular influences

The regulation of immune cell subsets homeostasis may be common to all cell subsets, or unique per population or lineage, based on functionality and developmental constraints. We noted that the number of associated hits was not uniformly distributed across immune cell subsets (**Figure 2**A, **Table S7**). This imbalance in trait-gene associations may be attributed to differences in regulatory complexity yet is most likely due to multiple complex issues involving both biological and technical limitations in experimental design. To overcome these issues and increase the strength of detected associations, we repeated our analysis in a second independently collected cohort of 24, non-overlapping, mouse CC strains, whose bone marrow we profiled by CyTOF similarly to the first cohort (see Methods, **Tables S8-S9**). This yielded successful validation of 788 genes (51%) from the initially detected set (**Figure 2**B, **Table S10** for adjusted *p*-*values*). Of interest, for 45 genes out of this set we noted that their mutation is known to affect leukocyte levels according to the Mammalian Phenotype Ontology (p<10^-12^ by *hypergeometric test*) (Smith and Eppig, 2009). For example, genes specific to the immune system: *Pax5*, *Cd28*, *Ltbp1* and genes with more housekeeping functionality: *Myo3b*, *Hnrnpr* and others (for full list see **Table S11**). Overlaying the genetic association by cell-type, we observed regulation that was overwhelmingly cell-type specific rather than shared across multiple cell types (**Figure 2**C). Per cell-type, we identified that the associated genes formed a functionally connected network, suggesting they act synergistically to control a cell subset’s frequency (**Figure S2**A-B).

**Figure 2.**
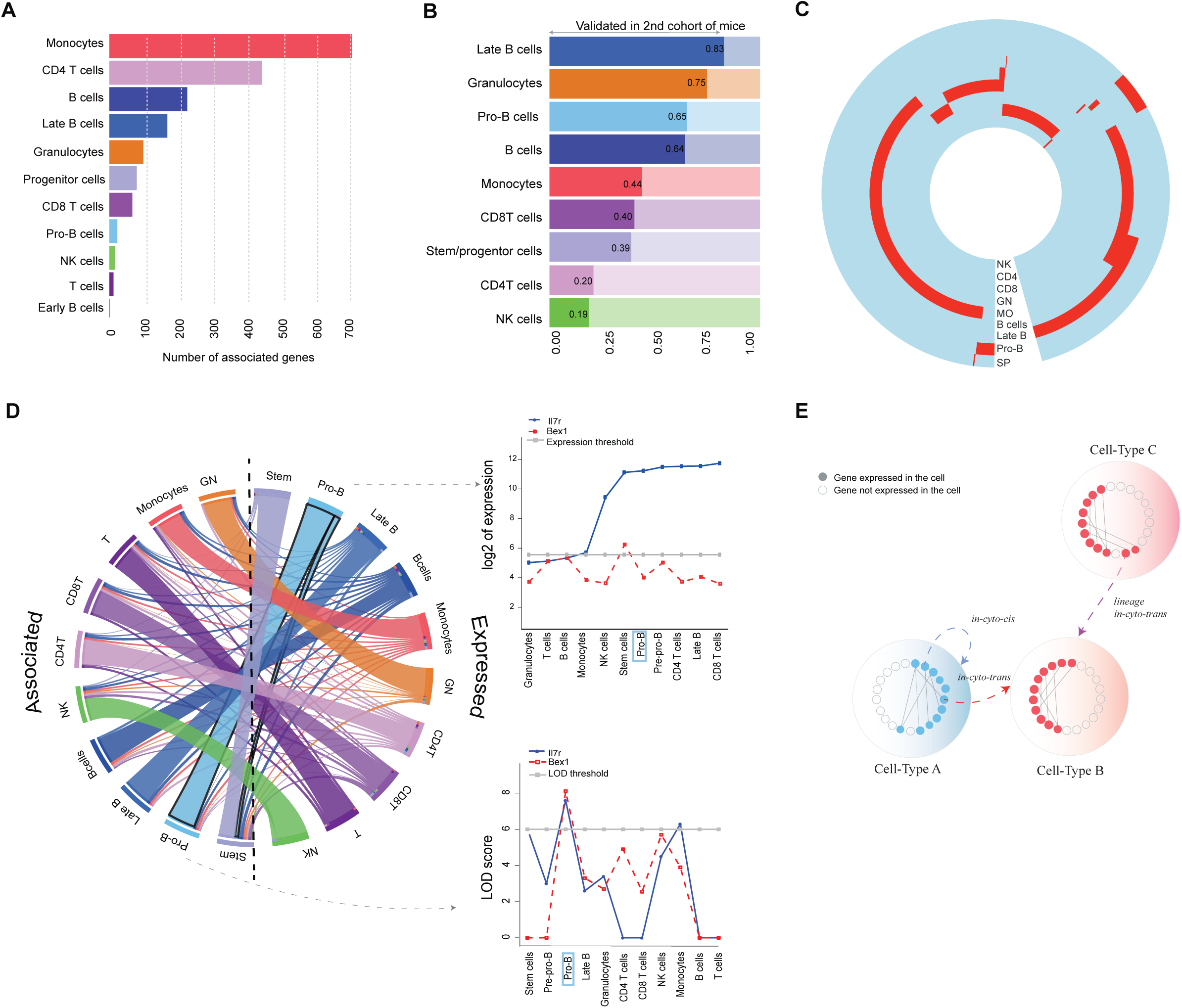
The regulation of immune cell subsets is cell specific and includes both *in*-*cyto*-*cis* and *in*-*cyto*-*trans* regulatory genes. **(A)** An imbalance in genetic associations across immune cell subsets. Bar-plot showing the number of associations per cell type. Bars are colored according to the cell type. **(B)** Validation of gene-trait associations in a second cohort of 24 non-overlapping CC mice strains (p<0.05 following BH adjustment). Filled-opaque bars show the percent of validation success for each immune cell subset out of the total number of associated loci for that subset. **(C)** A circular heatmap of per cell-type associated genes, abbreviation of cell types is indicated on each ring. NK-NK cells, CD4-CD4 T cells, CD8-CD8 T cells, GN-Granulocytes, MO-monocytes, SP-progenitor cell population. **(D)** Homeostatic condition of immune cells is predominantly determined by genetic variants expressed in the same cell subset. Split Circos plot, depicting the fraction of genes expressed in each cell subset (right) based on ImmGen sorted cell gene expression patterns and the cells in which we detected an association of a gene’s genetic variant with the cell’s frequency (left). The color of the ribbon is set according to the cell subset the genes are associated with. Inset (top) shows log2 expression across cell-types of Il7r and Bex1 and inset (bottom) shows the strength of association for Il7r and Bex1 across cell subsets (blue and red lines respectively, expression threshold in grey). Il7r is associated with pro-B cell subset frequency and also expressed in it, while Bex1 is associated with pro-B cell subset however is not expressed in it. (Short abbreviations used for: T-T cells, NK-NK cells, GN-Granulocytes, Stem-Stem and progenitor cells). **(E)** An illustration of regulatory process: expression pattern differs by cell-type (filled bubbles are genes expressed in the cell-type, hollow bubbles are not expressed). Two main patterns of regulatory genes: *in*-*cyto*-*cis* genes are expressed in and associated with the same cell-type (cell-type A regulates cell-type B blue line), *in*-*cyto*-*trans* genes are expressed by other cell-type. The latter type of regulation may exist between same lineage members (cell-type C regulates cell-type B purple line) or between members of different lineages (cell-type A regulates cell-type B red line).

Genetic regulation of a cell’s homeostatic condition may be determined by the cell subset itself, or by other cell subsets. For the majority (84%) of genetic variants we detected an association only in a single cell subset. To delineate the relationship between a gene’s cellular genetic association and its expression pattern we checked each gene’s cellular expression profile across the ImmGen sorted cell gene expression compendium (Heng and Painter, 2008) (see Methods). For the overwhelming majority of the genes whose variants were associated with a specific cell subset’s frequency we detected their expression to be specific for that same cell subset (from hereon *in*-*cyto*-*cis*), (**Figure 2**D, mean 86.5%, ranging between 83-100% across subsets). Whereas, 1.5% of genes were expressed in the same lineage, but not in the associated cell subset, and an additional 5% not expressed by any of the immune subsets whose association we tested. The remaining 12% we defined as *in*-*cyto*-*trans*, that is genes whose variants were associated with a specific cell subset frequency, but not expressed in that cell or in other subsets of the same lineage. The distribution of genes between these latter three genes sets was expression threshold dependent, yet their existence could not be discounted and robust to false discovery assessments (see Methods). Taken together this suggests that homeostatic composition is a polygenic trait which at the highest significance level is cell type specific, that is, determined for each cell independently by different genes; yet influences on each cell’s abundance by genetic variants of genes expressed in other cell-types may exist.

### Immune cellular profiles are governed by a polygenic network-of-networks with distinct architectural attributes

Given the complexity of the immune system we hypothesized that shared regulation of cell subsets is more pervasive but could not be detected due to stringent GWAS cut-offs. To overcome this, we leveraged the entire genetic association profiles, irrespective of association strength, of each of the 788 validated genes across cell types to identify those genes likely to be associated with multiple cells. To do so, we clustered gene-phenotype association profiles and discovered 13 modules of cell-gene association profiles (see Methods). We identified 419 multi-cell-associated (MCA) genes, out of the 788 validated genes, which exhibited high LOD scores across two or more cell types (**Figure 3**A). We noted that genes associated with the progenitor cell population (from hereon progenitor MCA) were enriched for MCA and associated on average with more cell-types than non-progenitor associated (**Figure S3**A, p<10^-42^ by *hypergeometric test* for MCA enrichment). Fitting with this, we noted that LOD scores of genes with the cell-populations below the association cutoff, were significantly higher for progenitor MCA genes than those LOD scores we observed for non-progenitor MCA genes (**Figure 3**B, p< 10^-6^ by *Wilcoxon*). Summarizing the results of this signal-propagation methodology yielded a four level grading of associated genes: first, a global regulatory group of progenitor MCA genes (178 genes), second, pan-lineage MCA genes common to two or more lineages (169 genes), third, lineage-specific MCA genes shared solely between cell types in same lineage (72 genes) and finally those genes associated exclusively with a single cell-type tested (369 genes).

**Figure 3.**
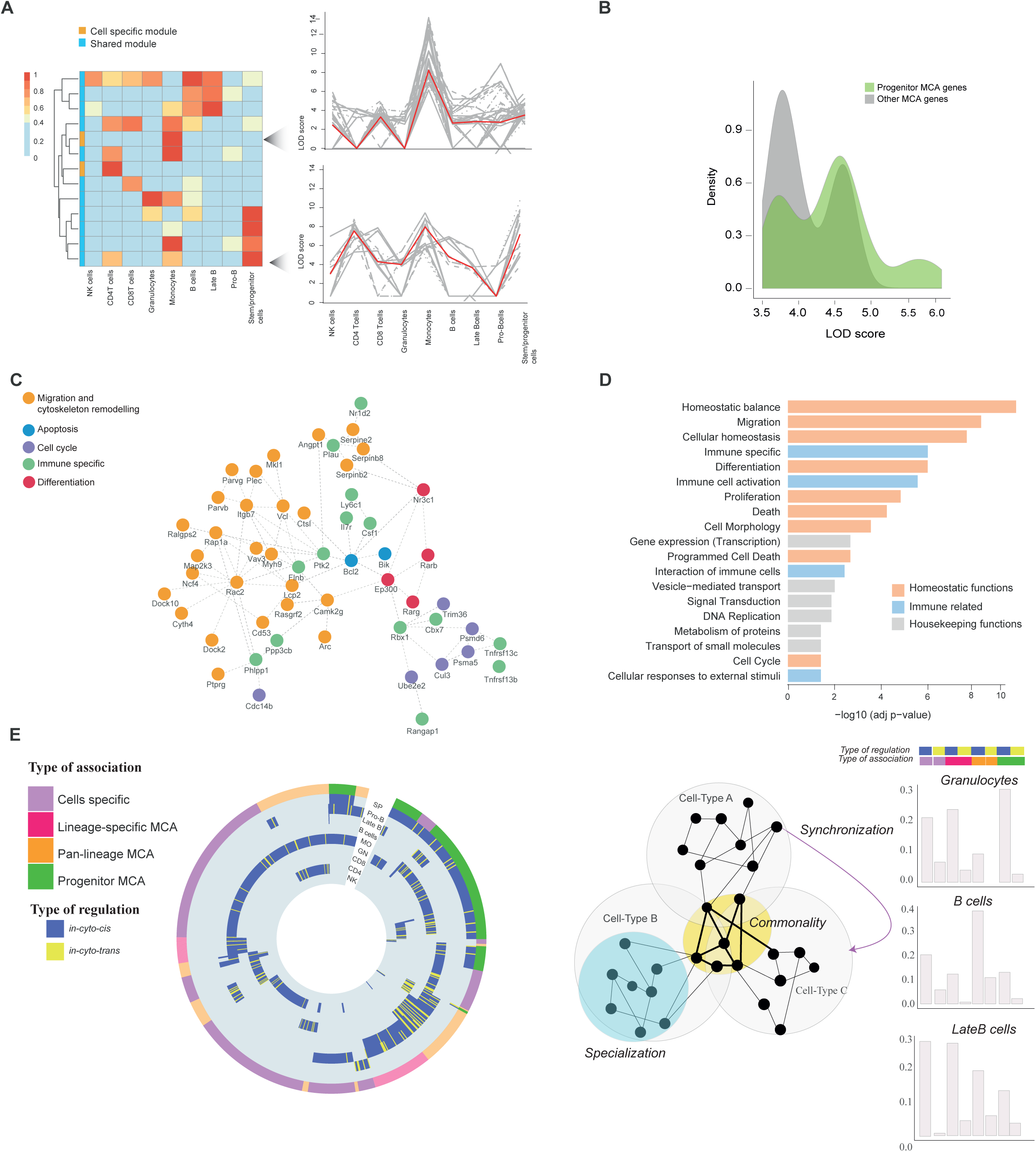
Polygenic network-of-networks regulates immune cell subset frequencies in homeostasis. **(A)** Signal propagation approach reveals additional regulatory layers of the immune homeostasis. Clustering gene association profiles across multiple cell types identifies modules of genes with similar association pattern, the percent out of maximal association median LOD score is shown for a module. Inset: individual module profiles, each line represents a gene in a module, red line shows median association for each LOD score. Modules are divided to cell type specific or shared across multiple cell types (blue or orange color in annotation bar respectively). **(B)** Progenitor MCA genes form a cell-gene association hub. Density plot of LOD scores of progenitor MCA genes (green) versus other MCA genes (grey) achieved in non-associated cell populations (p<10^-42^ by *hypergeometric test*). **(C)** Protein-protein interaction network of progenitor-MCA group of genes, the genes are colored according to their related pathways. **(D)** Immune cell subset abundance is largely determined by genetic variants of genes associated with homeostatic functionalities. Bar-plot shows functionally enriched categories. X-axis indicates -log10 adjusted p-value of the enrichment analysis (BH corrected Fisher’s exact test). Homeostatic functions indicated in orange, immune-related in blue and housekeeping in grey. **(E)** From left to the right: (i) Per cell type circular heatmap of associated genes following the signal propagation approach. Genes are colored according to regulation type (*in*-*cyto*-*cis* in blue, *in*-*cyto*-*trans* in yellow). Outer circle color reflecting type of association: cell specific, lineage specific MCA, pan-lineage MCA and progenitor MCA genes colored (purple, pink, orange, green accordingly). (ii) Immune regulation is performed by polygenic network-of-networks with distinct architectural attributes. *Commonality* dictated by shared genes, *Specialization* by cell-specific genes, *Synchronization* by genes affecting a cell-type they do not expressed in. (iii) Representative bar-plots for granulocytes, B cells and Late-B cells regulatory composition. Two color bars are showing the annotation of each group of genes, for example, first group has both blue and purple annotation, indicating cell specific *in*-*cyto*-*cis* group of genes.

From a functional perspective progenitor MCA genes formed a protein-protein interaction network (**Figure 3**C) whose members play a role in the regulation of cytoskeleton remodelling, migration (e.g. Rac2, DOCK2, FAK (Ptk2), integrin β7), apoptosis and hematopoiesis (Bcl2, Csf1, Il7-receptor, BLK); processes underlying key homeostatic functionalities whose dysfunction has been reported to drive immune-deficiencies and provoke cancer (Puel *et al.*, 1998; Ambruso *et al.*, 2000; Neri *et al.*, 2011; Schuetz *et al.*, 2011; Infusino and Jacobson, 2012; Dobbs *et al.*, 2015; Dezorella *et al.*, 2016). Generalizing on these functional observations, we performed a pathway enrichment analysis on all homeostatic network associated genes (**Figure 3**D, **Table S12** for significantly enriched functions, threshold at p<0.01 BH adjustment). We observed that 52% of functionally enriched genes were annotated for homeostatic functions and mainly were being divided into one of four processes: proliferation, cell death, differentiation and cellular movement (p =10^-8^, 10^-5^, 10^-4^, 10^-10^ respectively, BH corrected). These functions were observed in each cell subset (**Figure S3**B). Thus, that akin to mRNA or protein species abundance, cell subset abundance is largely determined by genetic variants of genes associated with turnover.

Combining the results of our gene-cell association grading with the information on mRNA expression cellular expression profiles, protein-protein interactions and functional analyses we performed conferred three attributes to the immune cellular homeostasis network (**Figure 3**E): (1) Commonality across cell-types of cell-abundance-trait determination dictated by MCA genes, (2) Specialization of cell type specific abundance dictated by *in*-*cyto*-*cis* cell-specific genes, (3) Synchronization of systems-level balance between cell-subsets dictated by *in*-*cyto*-*trans* genes. Taken together our analysis suggests that homeostatic composition is governed by a polygenic “network of networks” with distinct architectural attributes and functionally enriched groups fitting the distributed and coordinated multi-cellular biology of the immune system.

### Genetic variants of evolutionary conserved turnover genes dictate immune cell subset abundance

The architecture of immune homeostatic network and the central attributes which it confers may have risen over evolutionary timescales as the immune system developed to the rich multi-cellular communicating network it is today; beginning with an ancestral cell-type, the system diversified to include specialized cell-types which communicate with one another to achieve immune protection. Attributes such as shared regulation, a specialized cell-specific programs and inter-cellular synchronization program each pose a leap in the evolution of a complex system. We hypothesized that information on the evolution of these network attributes could be garnered through the evolution and function of genes tethered to each attribute. To do so, we used a gene sequence conservation score computed across 60 vertebrate species spanning an evolutionary time span exceeding 601 million years (see Methods). Testing differences in evolutionary conservation based on the expression of the functionally enriched genes, we observed that *in*-*cyto*-*trans* genes showed significantly reduced conservation score than *in*-*cyto*-*cis* genes (**Figure 4**A, p< 0.0002 by *KS test*), suggesting that synchronization functionality between cell-types evolved at later stages of the homeostatic network. However, analyzing differences in evolutionary conservation, we observed significantly higher conversation score of progenitor MCA genes compared to the cell-specific genes (**Figure 4**B, p< 10^-5^ by *Student’s t-test*), suggesting genes associated with cell cycle regulation in multiple cell-types, and in particular progenitor MCA, appeared earlier in the evolution of the immune system compared to genes with similar annotation but with a cell-specific association. Interestingly, we observed an enrichment of *in*-*cyto*-*trans* genes for MCA (p<10^-5^ by *hypergeometric test*), suggesting that MCA status may be achieved either via conservation of ancestral association (e.g. progenitor MCA) or by trans-regulation.

**Figure 4.**
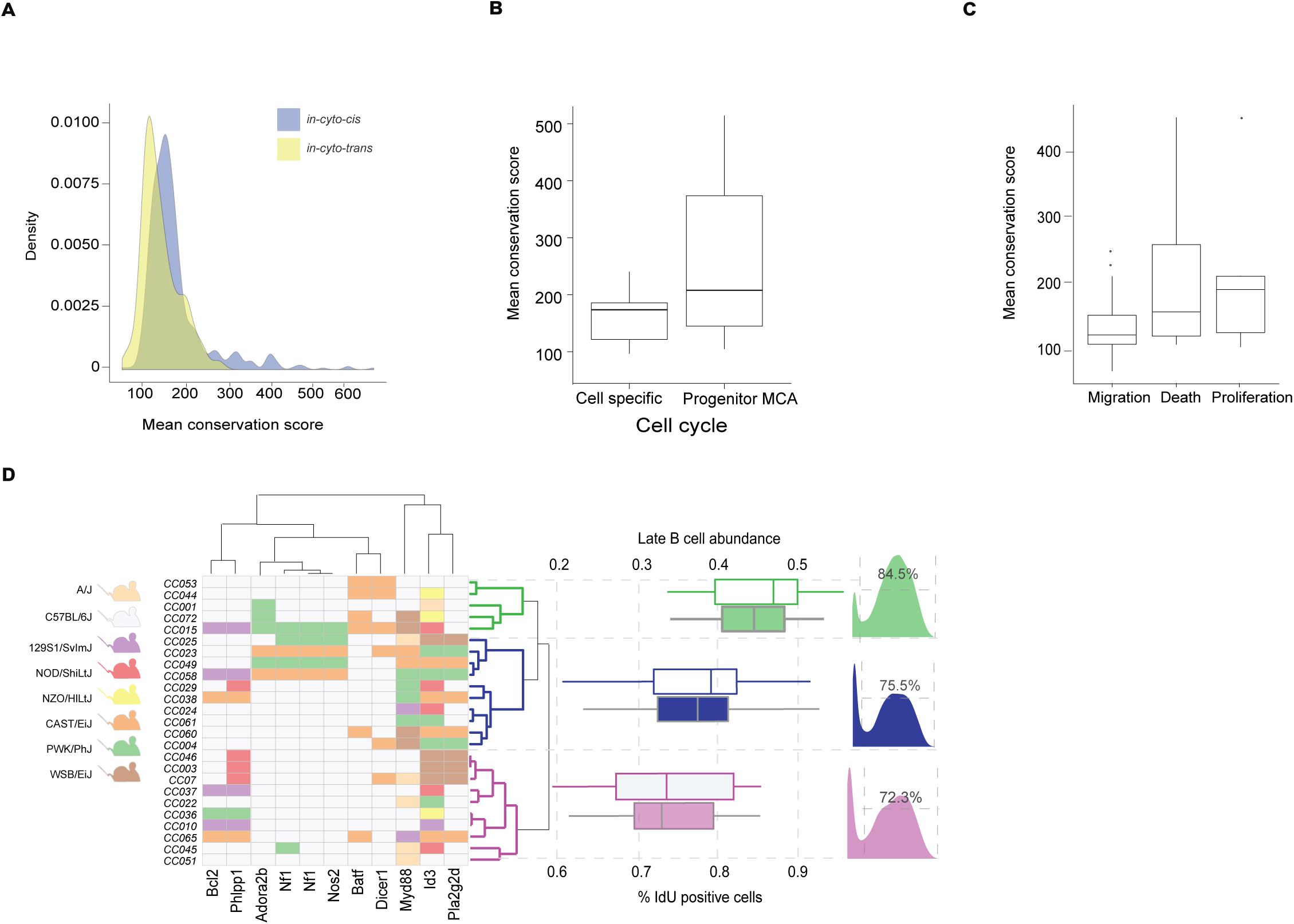
Immune cell subset abundance is determined by evolutionary conserved turnover genes. **(A)** Inter-cellular synchronization of immune cell homeostatic control evolved at later evolutionary stages. Density plots of mean conservation score for *in*-*cyto*-*cis* and *in*-*cyto*-*trans* group of genes (p< 0.0002 by *KS test*). **(B)** Box plots showing progenitor MCA genes associated with cell cycle pathway are more conserved compared to cell-specific genes with similar annotation (p< 10^-5^ by *KS test*). **(C)** Comparison of mean conservation score between enriched homeostatic functions groups of genes (p < 0.0005 by *ANOVA*). **(D)** Genetic similarity dictates phenotypic similarity. Clustered genetic profile of CC mice based on late B-cell subset associated proliferation genes. Alleles that differ from the C57BL/6J mouse are colored by founder strain. CC mice clustered by the hamming distance. (ii) For each clustered group, two boxplots are shown: (1) the abundance (colored contour, p<0.01, by *ANOVA*) and (2) percent of proliferating cells (colored fill, p< 0.05, by *ANOVA*), (iii) Histogram of IdU staining of a representative sample for each clustered group.

To show the importance of these turnover genes with trait determination, we sought to assess whether similarity in genetic profiles would yield a similarity in cellular profiles. Breaking down homeostatic associated genes by function, we noted proliferation genes showed significantly higher conservation score (**Figure 4C**, p< 0.0005 by *ANOVA*). We chose to focus on proliferation, given our evolutionary conservation analysis and that we could experimentally assess proliferation rate by measuring the percent of cells in S phase by IdU incorporation (Methods). We clustered mice from the validation cohort based on their genetic variants of validated proliferation genes associated with a specific cell-type (**Figure 4**D, left) and contrasted the differences in cell abundance and percent of proliferating cells in each subset between clusters (**Figure 4**D for late B-cells, **Figure S4** for monocytes, **Table S13** for proliferation estimates). We observed that the mice having PWK/PhJ or CAST/Eij founder strains contributing to the allelic variants of genes associated with late B-cell subset, had significantly higher proliferation rates (p< 0.05, by *ANOVA*) and higher subset abundance (p<0.01, by *ANOVA*). Taken together, these findings support a model of immune homeostatic network evolution in which a polygenic profiles similarity in these genes suffices to determine cell subset turnover and its abundance.

### Cell subset abundance determining genes are conserved in humans yet expand as a function of the environment

The immune system is affected by the environment which alters over an individual’s life history. Trait determination in such a scenario may result in genes being added or possibly even removed from the gene regulatory network governing the trait, as a function of whether a certain genetic variant plays a role in the biology condition at hand. Given the evolutionary conservation we observed for the genes governing the homeostatic network in the mouse, we next aimed to check how this network will manifest in environmentally experienced human adults. To do so, we analyzed the outcomes of three human GWAS studies aimed at identifying genes associated with immune homeostatic conditions. In total, these three studies tested the association over 300 traits of immune cell subset frequencies or counts, identifying total of 59 genes associated with one of these phenotypes. For these genes we observed functional enrichment that broadly split into two broad categories (**Table S14**): immune specific-related functionalities of response (e.g. ‘stimulation of lymphocyte’, ‘activation of lymphocytes’) which formed the overwhelming majority of functional enrichments, and a minority of functions annotated for homeostatic control. Beyond this, we noted were annotated for various immune diseases including influenza, celiac, rheumatoid arthritis (**Table S15**). Contrasting this functional annotation information with that we obtained in the mouse strongly highlighted the opposite trend, namely homeostatic control genes formed the majority of associated genes and enrichment for immune-specific-related functionality was predominantly absent. (**Figure 5**A left for mouse, right for human). The differences in lifespan and environmental exposure between mouse and man, suggests that genes associated with controlling immune cell abundance may be divided into genes whose functions are associated with environmental-interaction, such as TLR family members responsible for the alteration of the cell subsets due to pathogenic response, and those associated with homeostatic control independent from the environment, such as ATG family members responsible for autophagy.

**Figure 5.**
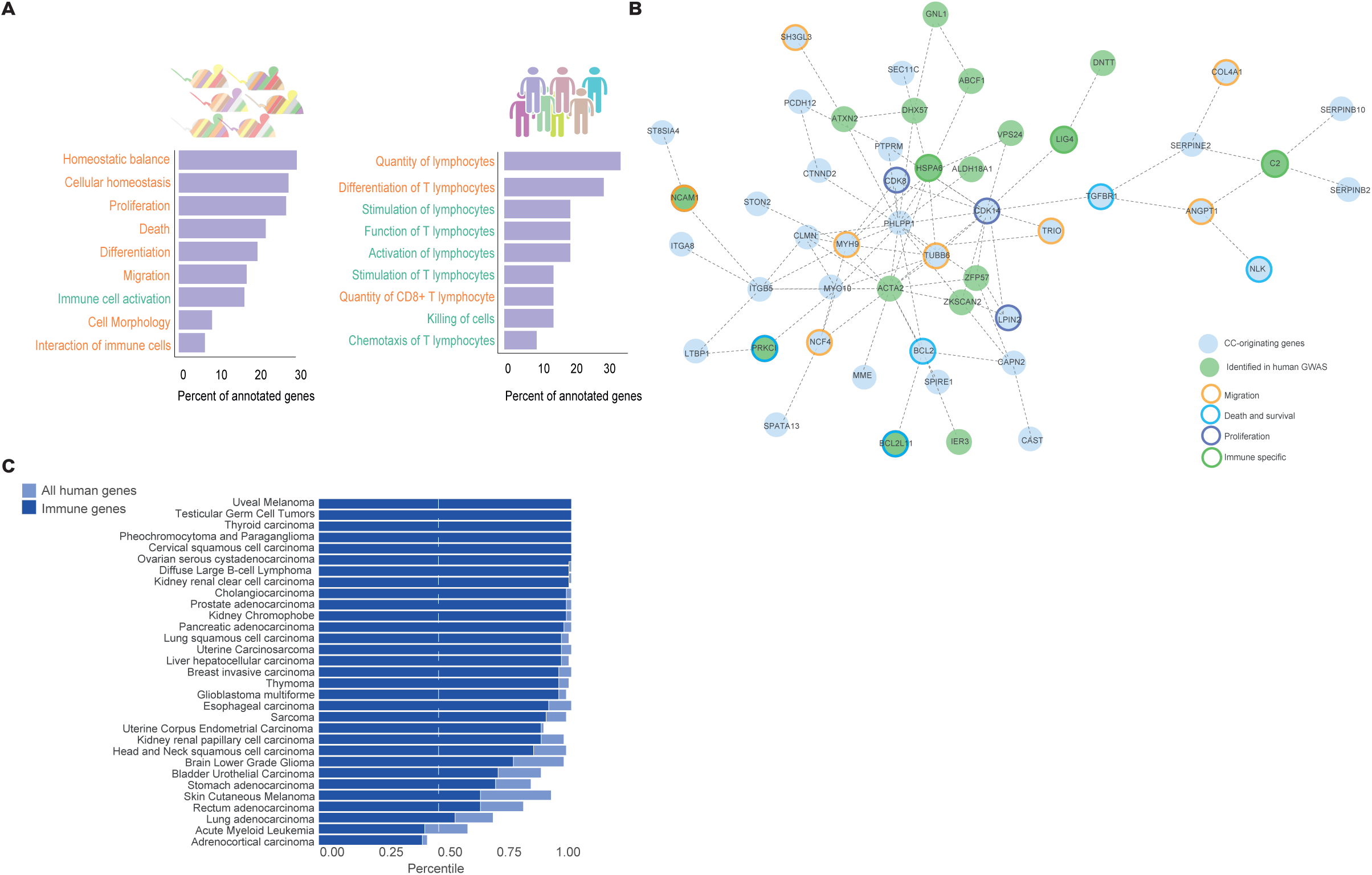
Cell subset associated polygenic network validates in human GWAS and expands during lifetime as a function of environmental exposure. **(A)** Human and mouse associated genes show opposite trends in functional annotations. Enrichment analysis of genes found in CC mice study and three human GWAS categories with the highest number of genes are shown in the bar plots. The names of the categories are annotated in orange for homeostatic functions, green-functions that are related to immune response. **(B)** Homeostatic control genes identified in the CC and validated in human and the human set from GWAS formed a network of 51 genes with direct interactions between immune response and homeostatic genes. Genes are colored by homeostatic functions (migration, death and survival, proliferation and immune response colored respectively orange, blue, purple and green), no coloring-genes with no enriched functions. **(C)** Tumors may preferentially mutate genes from the homeostatic control network. Genes associated with immune cell subset frequency are at the top percentile of mutated genes in cancers, compared to equally sized random controls. Bar-plot shows different TCGA cancers with percentile calculated against a background consisting of genes expressed in immune cells (blue), or all human genes (light blue).

We reasoned that the network governing an organisms’ cell subset abundance must include genes whose function is directly associated with cellular turnover. These we predominantly identified only in the mouse but not the human. We analyzed the association of validated mouse orthologs with one or more of 130 measured immune cell subset traits in a human immunophenotyping GWAS (see Methods) (Roederer *et al.*, 2015). Testing 42,616 relevant SNPs, corresponding to 710 orthologous genes, we detected 47 genes which were associated with the abundance of one or more cell subsets (**Table S16**, p< 1.9^∗^10^-6^ using an adjusted significance threshold, see Methods). Next, we asked whether the 47 CC-originating genes interact with one another and to those associations discovered by the human GWASs. Protein-protein interaction analysis of the human associated genes pooled from all three GWAS, did not form a coherent network of interacting genes (**Figure S5**A). Yet, incorporating the CC-originating genes with those identified in human, we observed a large connected component consisting of 51 genes with direct interactions between immune specific and homoeostatic control genes (**Figure 5**B, **Figure S5**B, see Methods). In agreement with the human functional enrichment analysis, the function of genes in the connected component was not equally spread between species; rather immune specific functions were derived from the human studies and homeostatic from the CC mice. Considering the complementary nature of the mouse and human hits, and the differences in environment between the two species. Taken together, this suggests that over time as a function of environmental exposure the gene regulatory network governing immune cell abundances changes and adds additional environmentally sensitive components, with genetic variants of environment-interacting genes regulating cellular abundance via their interaction with environment-independent genes.

Given the homeostatic functionality we detected in the associated genes, we reasoned that their mutation may play a role in diseases in which the normal process of proliferation and/or death is disrupted. We thus leveraged information regarding patient somatic mutations in TCGA cancers, including both solid and blood tumors, to check the behavior of genes associated with control of cell subset frequencies (see Methods). Across the majority of cancers, we observed a strong enrichment for mutation accumulation in this gene set compared to equally sized random controls sampled from the entire human genome as well as to a more conservative background consisting of genes expressed in immune cells only (p< 10^-7^, p<10^-17^ respectively by *Fisher’s combined probability test*, **Figure 5C** when performed for 106 cell subset associated genes, **Figure S5**C when restricted to 47 CC derived genes validated in human, see Methods). This mutational enrichment was observed even following exclusion of known proliferation and death annotated genes (p<10^-6^, p<10^-7^ respectively by *Fisher’s combined probability test*). Taken together this suggests that tumors may preferentially mutate member genes of this homeostatic control network for co-option, from their original immune origin, to control growth.

## Discussion

Here we coupled the high variability of Collaborative Cross mice with the high dimensional phenotypic capabilities of mass cytometry to identify a global genetic architecture that is responsible for between individuals’ homeostatic differences. Our analysis revealed an unprecedented number of novel associations of genes with immune cellular homeostasis. Using a network propagation approach, we detected a complex gene regulatory network made up of distinct cell-type networks and an inter-cellular network connecting them which governs homeostatic balance. Moreover, this genetic network is enriched for homeostatic functionalities, whose genetic variants are the origin of individual immune variation and can determine phenotype as we experimentally show. By hypothesis driven testing we validate a subset of the network in human GWAS which exhibits mutational enrichment across multiple cancer types.

Control of cell abundance is a key feature of all immune cells in response to stimuli. The evolutionary sequence conservation we observe for homeostatic network genes mirrors a model of evolution whereby the immune protection shifted from being the responsibility of a single cell types, to multiple cell types which work in a synchronized manner to achieve an emergent immune protection. Progenitor MCA genes, are oldest and may have formed the earliest part of the network, with cell-type specific genetic variants following as cells could tune their abundance by specialized mechanisms fitting their biology. Finally, genes with *in*-*cyto*-*trans* pattern of regulation could develop for coordination of inter-cellular homeostatic balance (**Figure 6**A). More broadly, this immune system growth model would suggest that over evolutionary time scales, as the system becomes more complex the number of genes involved in homeostatic determination would grow (**Figure 6**A, top, ‘black line’).

**Figure 6.**
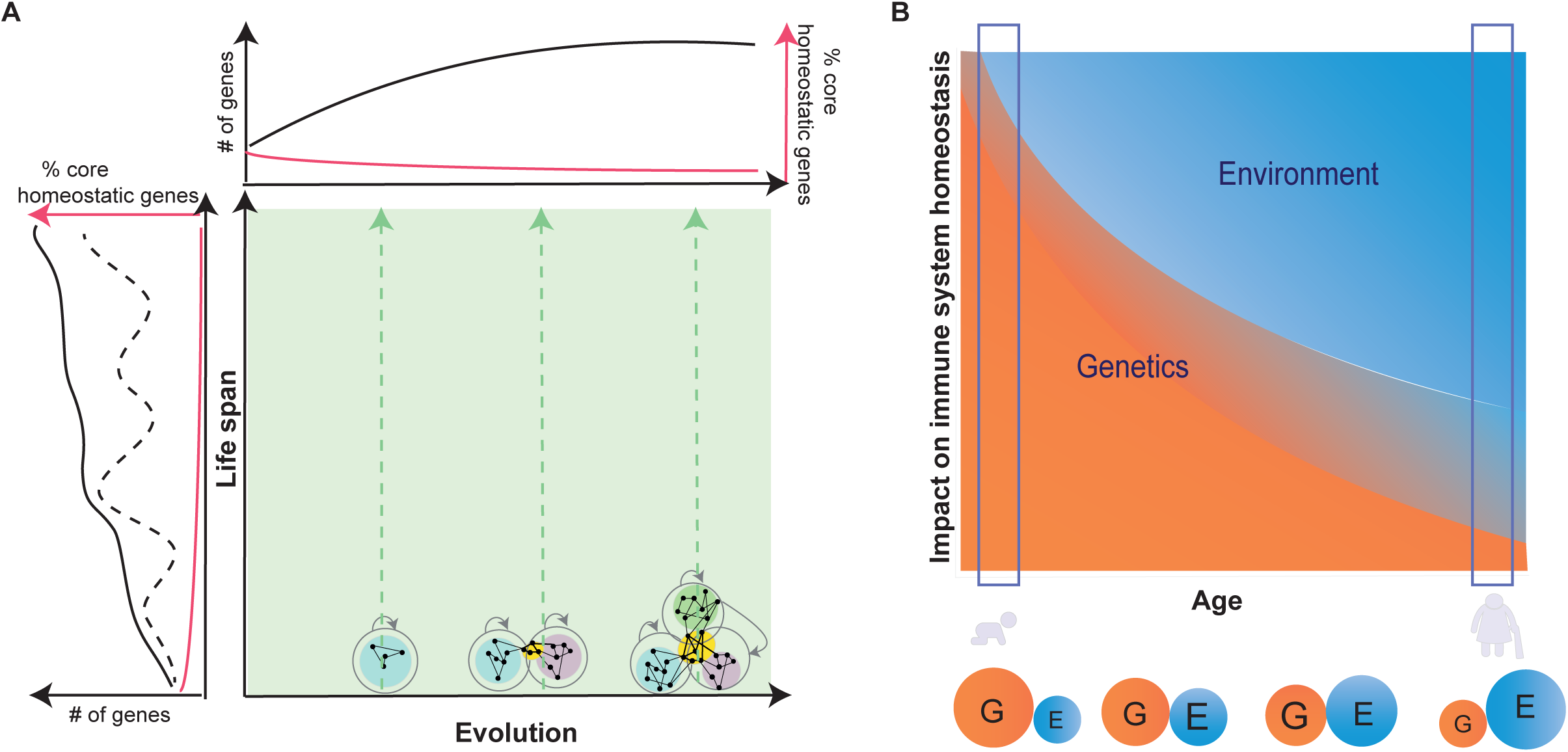
A model of a multi-cellular regulatory network dynamics as it alters as a function of system evolution and environment. A. Complex trceaits are determined by genetic regulatory networks that change over both evolutionary and life span time scales. On the evolutionary axis, when only a single cell type existed, a genetic network which regulated that cell type included mainly MCA genes. With time the system diversified to include additional cell types, such that more cell specific genes were added, finally genes responsible for synchronization were created to control the abundance of multiple cell types. On the life span axis, this polygenic network changes with life span, as with increased environmental exposure additional genetic variants begin playing a role in the regulatory network. The number of core homeostatic genes, set of genes directly associated with the phenotype, those responsible for homeostasis maintenance and the regulatory network size change over individual’s life span and possibly over evolutionary time scales.
B. Both genetics and environment impact the immune system and the impact of the latter grows with age. Identifying genes which underlie immune cellular homeostasis in early and late life stages may thus result in non-overlapping gene sets. This can be generalized to all environmentally sensitive systems where both genetics and environment impact the phenotype, and where the balance between them shifts over time.

Beyond evolutionary timescales, our results suggest that the polygenic network which determines immune cellular homeostasis also changes on a much shorter time scale, that of an individual’s life history. Our analysis of human GWAS suggests that as environmental exposures occur through life, additional genes begin playing a role, possibly becoming the more dominant effectors of the network phenotypic outcome. The total size then of the gene regulatory network associated with the trait expands over lifespan, though the number of genes with a large effect size on the phenotypic outcome may vary over the course of an individual’s lifetime (**Figure 6**A left, ‘black solid and dashed line’, Figure **6**B).

Our study sheds light on several burning issues related to polygenic trait determination: First, and recently hotly debated issue (Boyle, Li and Pritchard, 2017; Wray *et al.*, 2018), is whether a set of ‘core’ genes exists in a polygenic network which are directly associated with phenotype functionality, and thus more important to investigate. Our choice to study cellular homoeostasis is illuminating in this regard, as based on mathematical principles defining turnover as a set of equations describing production and removal, a core set of genes may be reasonably well defined. Our results suggest that turnover genes, are the core of the network (52%) yet over an individual’s life history and possibly over evolutionary time scales, the relative contribution to the network of this core homeostatic gene set will shrink due to addition of other genes or substitution of the relevant core. This implies, that a focus on core genes, even in this well-defined trait may be problematic (**Figure 6A** top and left, ‘red line’, **Figure 6B**). We expect this to be even more problematic, when a trait’s biology is difficult to ascertain or may even change as a function of life or disease progression. A second interesting point is that 46% of associated genes in our study are associated only with the regulatory network of a single cell-type, whereas it has been suggested that the genetic contribution of cell-type specific regulatory elements is minor (Boyle, Li and Pritchard, 2017). This contrast may be explained by the increased cell-type level resolution of our study whereas the prior expression profiles of genes were obtained at a tissue level (‘The Genotype-Tissue Expression (GTEx) pilot analysis: Multitissue gene regulation in humans’, 2015). Last, due to power considerations we exclusively studied SNPs located in coding regions, despite the fact that it has been suggested that complex trait variation is predominantly determined by non-coding regions (Pickrell, 2014). Our discovery of a large number of genetic associations in coding SNPs does not discount the non-coding regions, but does suggest an underestimate for the role alteration in protein structure may create in a gene regulatory network.

The large variation exhibited by CC mice, coupled with their controlled genetics and clean environment make them an ideal model for systems level studies, providing opportunity usually unavailable in human – such as to profile bone marrow. It would be interesting to analyze mice under controlled simulation of real-life environmental exposure, as our inferences on changes in the homeostatic network as a function of environmental exposure currently come from studying humans. The polygenic network we uncovered in that the bone marrow might be reflected in other tissues as well, and likely affected by other cell types in the tissue. It might be of interest to understand how the polygenic network that governs immune cell subset abundance communicates with other regulatory networks in the body, for example microbiome, which have been shown to have an impact on immune cell homeostatic balance (Snijders *et al.*, 2016).

Evidence of immune cellular profiles or immune states being of clinical relevance for diagnostic purposes has been shown across multiple disease conditions and health outcomes. These states are reached through a developmental trajectory which begins in early life. The position of an individual on this trajectory is a function of the interaction of the genetic network we identify here with the environment in which the individual resides. This suggests that our results may be built on for tailoring immune cellular clinical assays for personalized medicine.

## Acknowledgements

We would like to thank Fernando Pardo Manuel de Villena, Darla Miller and Ginger Shaw for providing us the CC mice strains and help with the mouse experiments. Asya Rolls, Tamar Ben-Shannan, Hilla Azulay-Debby, Ben Korin and Maya Schiller for help with mice work and manuscript revision. Amit Ziv-Kennet for help with data preprocessing and Fuad Iraqi for initial setup of the system. We thank John Tsang, Motti Choder, Ayelet Segre, Doron Melamed, and Gil Atzmon and members of the Shen-Orr lab for fruitful discussions. This study was supported through generous support of the Israeli Science Foundation (grant 1365/12), Applebaum foundation, MALAT and the Colleck Research Fund.

## Author Contributions

T.D., E.S, S.S.O designed the study, T.D., E.S., A.A, Y.O. performed the experiments; T.D, S.S.O, E.S., B.L, R.N., Y.A, M.G, performed the analysis; T.D., E.S, B.L, M.G, S.S.O wrote the manuscript.

## STAR Methods

### Samples collection and processing

*Mouse samples*-founder strain males (n=3 per strain) of the A/J, C57BL/6J, 129S1Sv/ImJ, NOD/ShiLtJ, NZO/H1LtJ, CAST/EiJ, PWK/PhJ, and WSB/EiJ strains were purchased from The Jackson Laboratory and sacrificed at 6-8 weeks. For the Collaborative Cross strains, the Systems Genetics Core Facility (University of North Carolina) provided 129 mice aged 8-14 weeks from 55 different complete lines, at least two mice per strain. Mice genetic information can be found at http://csbio.unc.edu/CCstatus/. Mice were bred and sacrificed at UNC facility. All procedures involving animals were performed according to the Guide for the Care and Use of Laboratory Animals with prior approval by the Institutional Animal Care and Use Committee within the Association for Assessment and Accreditation of Laboratory Animal Care-accredited program at the UNC at Chapel Hill (Animal Welfare Assurance Number: A-3410-01). Bone marrow was collected from necropsy following humane euthanasia by CO_2_. Bone marrow was flushed with cold CSM (PBS + 0.5% BSA) from the femur and tibia using a 27.5-gauge needle and a 10mL syringe to achieve a single cell suspension.

*Human samples*-bone marrow aspirates from AML patients presenting at the Hematology Department in Rambam Medical Health Care Campus were collected. All patients gave informed consent according to the declaration of Helsinki (IRB number: 0573-10). Mononuclear cells were separated by centrifugation over a layer of LymphoprepTM (Axis-Shield PoC AS, Oslo, Norway) and then stored in freezing medium [fetal bovine serum (FBS) with 10% DMSO] in a liquid nitrogen tank.

*Cell staining*-Primary conjugates of mass cytometry antibodies were prepared using the MaxPAR antibody conjugation kit (Fluidigm Inc.) according to the manufacturer protocol and optimal concentration was determined by titration. Cells from each sample were washed twice and a total of 3 million cells were used for staining. Cells were resuspended in 500ul containing 1:2000 Rh DNA intercelator for 20 min of live/dead cells staining. Samples were washed with CSM buffer and resuspended in total of 100 ul metal-tagged antibody mix for cell surface markers staining for 1 hour. Cells were then fixed in 1.6% PFA (Sigma-Aldrich) in a total volume of 200 ul and stored at 4^0^C. DNA intercalator Ir191/193 staining was performed post-PFA removal for 20 min at 1:2000 concentration in 500 ul volume. Finally, fixed samples were washed 3 times with DIW immediately prior to acquisition.

*Samples acquisition and data analysis* -Samples were acquired using CyTOF 1 machine at 500 events/sec for a total of 100-200K events. Internal metal isotope bead standards were added for sample normalization as described (Finck *et al.*, 2013) to account for the decline in mean marker intensity over time. Acquired data were uploaded to a Cytobank web server (Cytobank Inc.) for data processing and gating out of dead cells and normalization beads. To account for intra-run declines in mean marker intensity over time, we performed a within-sample-over-time normalization step by using a running window to adjust mean marker intensity throughout each individual run such that the mean expression over time was equal to that measured at the beginning of the run. We manually gated all the major cell populations in human and mouse bone marrow (**Figure S1**B-C). Resulting phenotypes were than exported and adjusted to the total number of cells in the sample. Cell subset frequencies for each sample are summarized in (**Table S4**).

### Phenotyping of immune variation and mapping of associated genetic loci

Single cell data of bone marrow populations from CC mice were analyzed by Citrus, a high dimensional clustering analysis algorithm. 40,000 cells were sampled from each sample and the minimum cluster size was set to 1%. The resulting tree structure organizes clusters of single cells sharing similar set of 20 markers. Clusters are organized in a hierarchy with leaves of the tree in the periphery being more specific and those inward aggregating multiple leaves. Next, we performed PCA analysis on clustered bone marrow cell subset frequency profiles of the 15 CC strains and 3 founder strains, which had a full set of phenotypic markers and calculated the Euclidian distance between each two mouse samples taken into account first two PCA dimensions. We repeated the analysis for ten human samples. Finally, we mapped manually gated populations to high dimensional clusters using Scaffold algorithm, which calculates distance between populations based on a predefined set of phenotypic markers (**Table S2** for the list of selected markers for mapping).

*Filtering of loci*- to reduce the number of multiple hypotheses tests, we chose to focus on associations with genomic loci that passed the following filter criteria: First, those loci which contained genes for which at least one of the CC founder strain is predicted to have an altered protein structure due to a sequence change compared to C57BL6 in exons of protein coding genes (Keane *et al.*, 2011; Yalcin *et al.*, 2011); Second, those genes determined to be expressed in immune cells based on Immgen consortium (Ericson *et al.*, 2008). Using these two filtering criteria reduced the number of tested loci from 77,725 to 15,470.

*Gene-Phenotype Association*- we performed QTL mapping using DOQTL R package and MegaMUGA SNP chip (https://csbio.unc.edu/CCstatus/index.py) given the CC genotype reconstruction and the probability of descent from one out of eight founder strains for each genomic interval. We searched for significant association between immune phenotype and each genomic locus, using an additive linear model. To check for stability of observed association we used a *leave*-*one*-*out* approach, calculating the strength of association. We only selected the associations which passed the threshold of significance in a strain independent manner. We set a significant association threshold based on the FDR of same population. We adjusted the threshold such that will allow an FDR of 5% and less. We calculated FDR for each phenotype separately by shuffling each time the mice ancestry in the locus and calculating the strength of the association, followed by a *leave*-*one*-*out* approach as previously described.

### Functional gene classification and validation of genetic associations

*Gene expression*- We estimated expression of genes according to ImmGen, using a threshold of log2(47) (Ericson *et al.*, 2008), as a threshold for genes of intermediate and high probability of expression. To be conservative towards calling *trans*-*associations* based on the expression of a gene in the associated cell-type or another cell-type, we considered intermediate and high threshold of expression genes as expressed in a cell.

*Validation of gene*-*trait association*- We used a second cohort of mice to validate our findings. Each significant gene-phenotype association we identified in the first cohort was tested for significance in the second cohort (p<0.05).

*Enrichment analysis*- we performed pathway enrichment analysis with IPA software (Ingenuity Pathway Analysis Qiagen), using the core analysis module. Repeating related functional terms groups were grouped together and the group with highest adjusted p-value and total number of genes for each functional enrichment is shown. Functionally enriched genes were annotated for at least one of four main processes: proliferation, cell death, differentiation and cellular movement.

*Validation of variants of proliferation genes*- We injected the mice i.p. with 50mg/kg body weight IdU (5-Iodo-2’-deoxyuridine) twice: 48 hours and 24 hours prior to euthanasia. To test for differences in IdU levels we manually gated all the subsets for IdU positive and negative cells. Mice were clustered, per cell-subset, by hierarchal clustering, based on the loci identified as associated with the cell-subset in the first cohort. One-way ANOVA was performed in order to test for difference in percent of proliferating cells (*p*<0.05).

### Genetic network analysis

*Signal propagation approach* - for each cell type we chose relevant modules by comparing the median association LOD scores. We considered a module as associated with a cell type if its median LOD score was at least 40% of the highest median calculated for that module. Next, we constructed a network including all cell types and associated genes, such that for each cell type, genes found by QTL mapping and genes from signal propagation procedure were included.

*MCA distribution comparison*- we identified genes associated with progenitor cell population (i.e. genes member in one of six modules associated with progenitor cells). For these genes, all of which were MCA genes based on the incorporated modules, we computed across all genes, the distribution of LOD scores in those cell-subsets for which their association was below the association threshold (i.e. those cells with which they were <u>not</u> associated). This distribution we compared with a similar one obtained for MCA genes which were not members of these six modules (i.e. the five other MCA modules).

### Human GWAS analysis

We obtained data from the study published by Roederer and colleagues (Roederer *et al.*, 2015). Per trait, files (151 such files) contain association p-values for all SNPs tested in the study. Using Biomart R package, for each gene of interest we estimated which SNPs are in its closest neighborhood, the SNP with lowest p-value was chosen to account for gene-trait strength of association. We tested all genes found in the first cohort of our study and were validated in the second cohort, adjusting the GWAS threshold to be 1.9^∗^10^-6^ according to the number of SNPs we tested and correcting for 129 traits (out of 151) of cell subset frequency in the GWAS study. In such a way we identified 47 genes which passed the threshold and were associated with 4 immune traits. To assess the false discovery rate of the procedure we calculated the FDR for each LOD score by 100 randomly choices of list of genes (**Figure S5**B). Next, we searched for experimentally validation of protein-protein interaction between 106 genes (59 identified directly in one of the three human GWAS and 47 we identified in CC and validated in human). We identified a large interaction network covering 51 genes (16 of which were previously detected in the human and 35 novel human findings, stemming from the mouse).

### Cancer mutation enrichment

We obtained somatic mutations of cancer patients TCGA. For each cancer type we computed the overlap between genes mutated in that cancer with a random gene set of equal size to the remaining set of identified genes post filtering for filtering for one-to-one orthologues. We performed this computation 100 times to generate a distribution, and then checked the percentile at which the identified set of genes placed. We considered two sampling options, to sample genes out of the whole human genome or only from genes that are expressed in immune cell subsets. To conduct a more conservative test, we checked whether the results remain stable even when taking out genes that were previously functionally annotated as proliferation and death genes by IPA and/or GO. As such, we analyzed the data in four conditions: every combination of background and gene set options.

**Figure S1.**
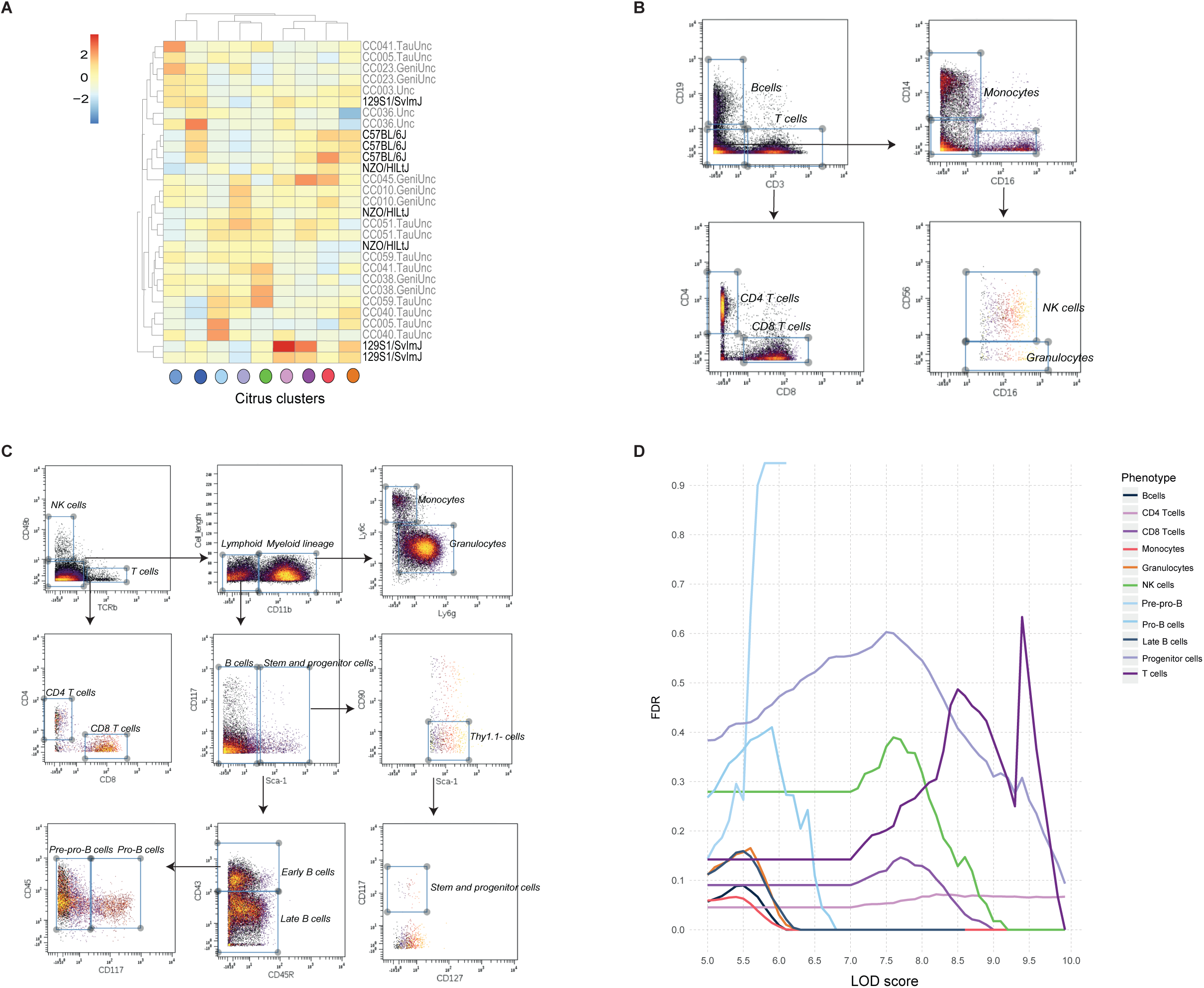
Gating strategies that were used for mouse and human bone marrow samples and calculated FDR for QTL mapping results, Related to Figure 1. A. Phenotypic profiling of bone marrow populations follows genotypic similarity. Heatmap of scale Citrus cluster frequency (columns) for CC mice and founder strains (rows). Founders indicated in black and the CC strains in grey.
B. Gating strategy of human bone marrow samples, 7 populations.
C. Gating strategy of mouse bone marrow samples, 12 populations.
D. Calculated FDR for each population is shown as a function of the LOD score.

**Figure S2.**
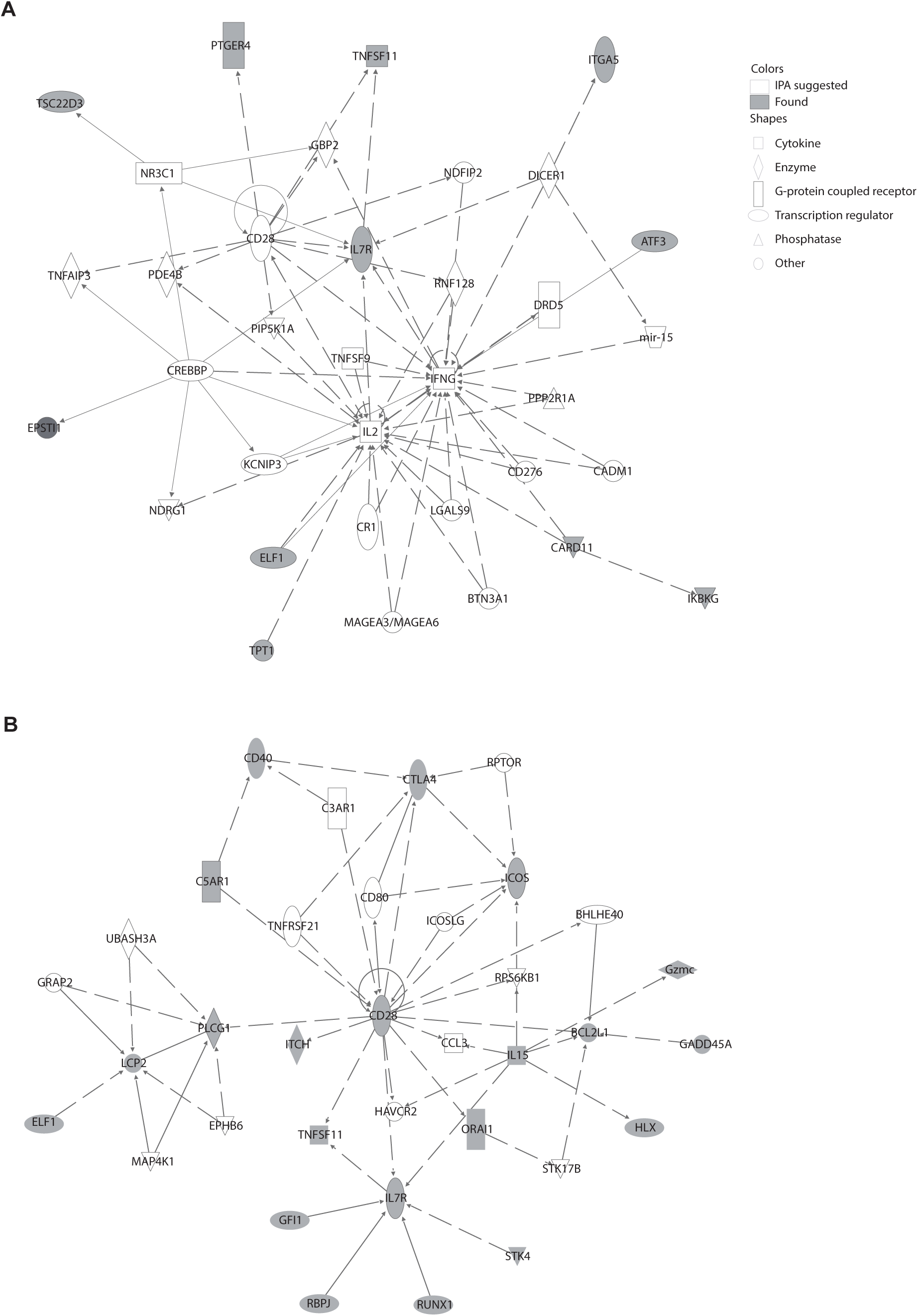
Cell specific network of regulation, Related to Figure 2. A. Pro-B cell specific network.
B. Monocyte specific network. Gene shapes are annotated based on IPA gene ontology, genes that were found as associated with monocyte and Pro-B cell subset frequencies are colored in grey.

**Figure S3.**
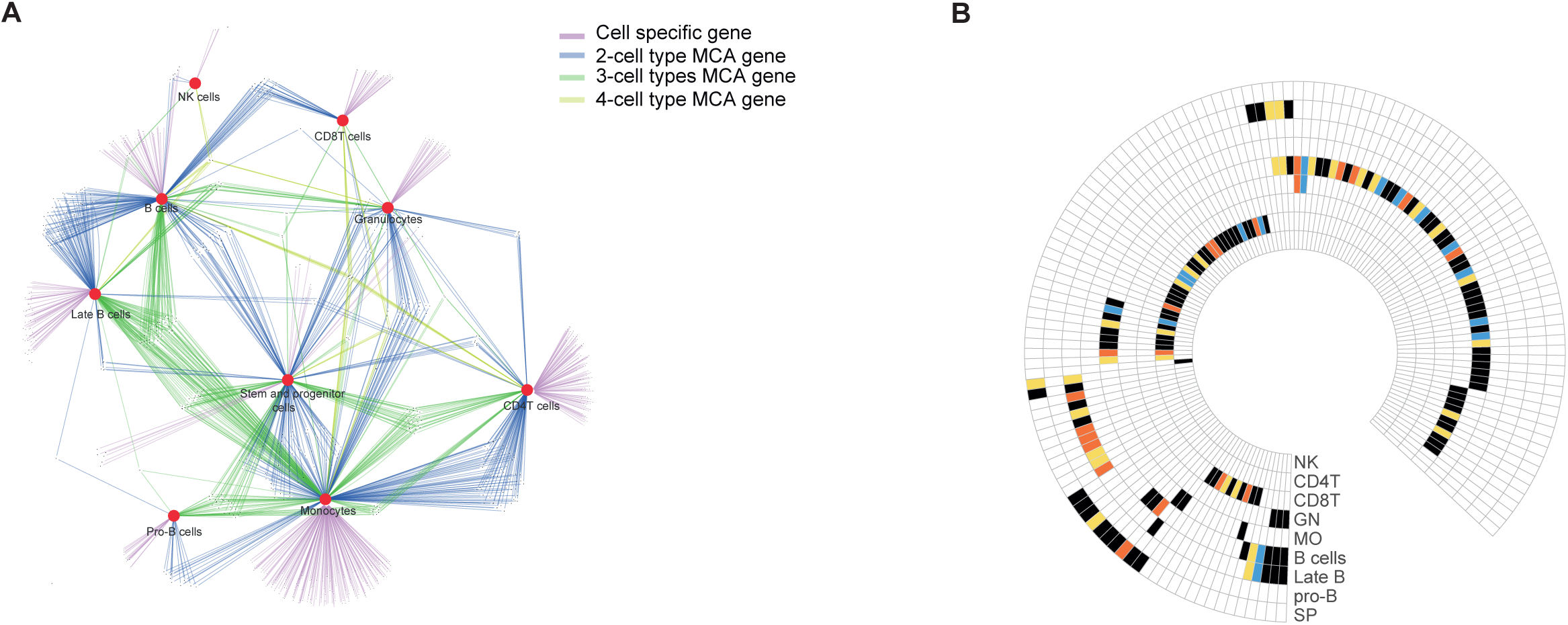
Polygenic network with different level attributes regulates immune cell subset abundance in homeostasis and has an enriched ‘core’ of turnover genes, Related to Figure 3. A. Genetic network of immune cell subset frequency associated genes, multi-cell-associated (MCA) genes are colored according to the number of cell types they associated with.
B. For every cell, genes associated with each of these four functions are found associated in a cell-specific manner. The subset identity is shown as: MO-monocytes, CD4T-CD4 T cells, GN-Granulocytes, Late B-cells, SP-Stem and progenitor cells, B cells, CD8T-CD8 T cells, NK-NK cells, pro-B cell. The colors of the enriched functions: blue-proliferation, orange-death, yellow migration, black-multi functional gene (all differentiation genes are multi-functional).

**Figure S4.**
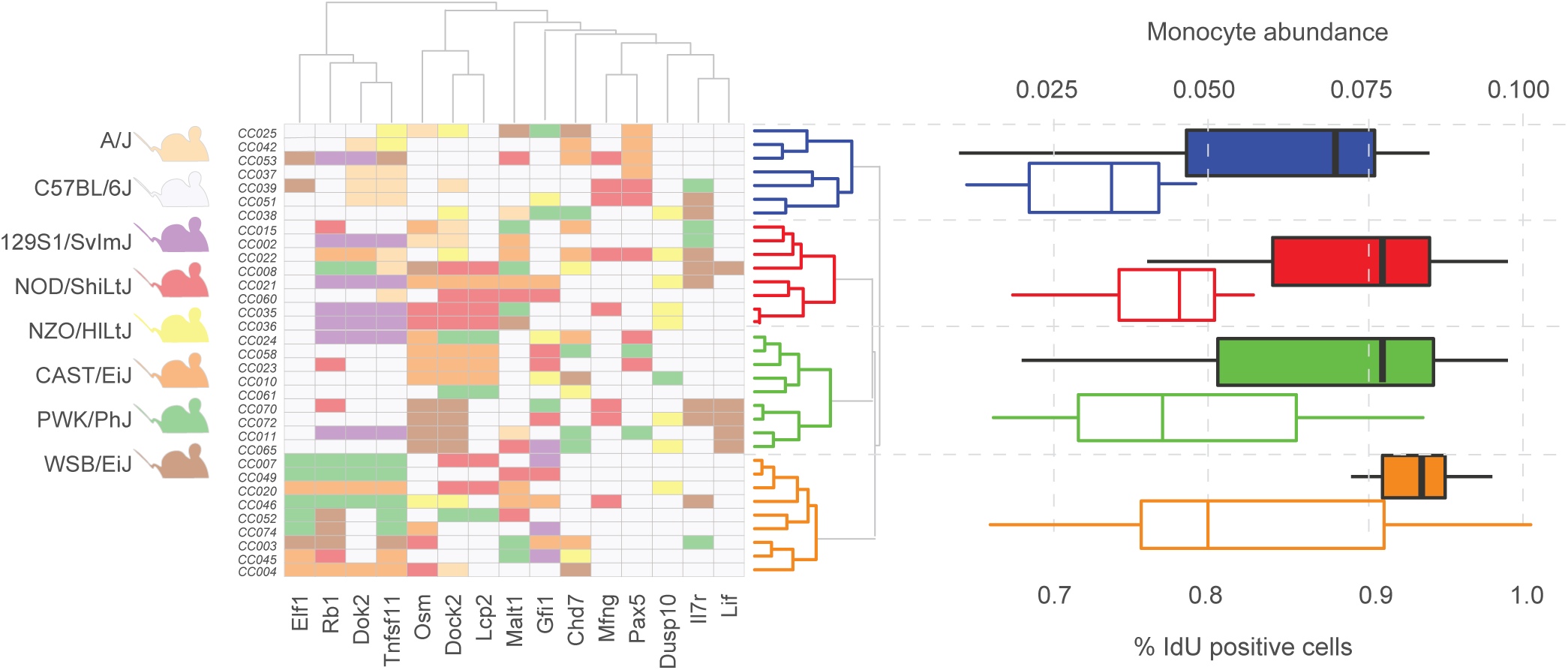
Validation of the proliferation function, Related to Figure 4. From the left to the right: (i) Clustered genetic profile of genes that are associated with monocyte cell subset frequency for each mouse. All possible alleles that differ from the C57/BL6J mouse allele for each gene were extracted, and the resulting profile across all associated genes was clustered by the hamming distance. (ii) For each clustered group, two boxplots are shown: (1) boxplot with colored contour for the abundance and (2) with a color fill for the percent of proliferating cells (bottom and top axes respectively).

**Figure S5.**
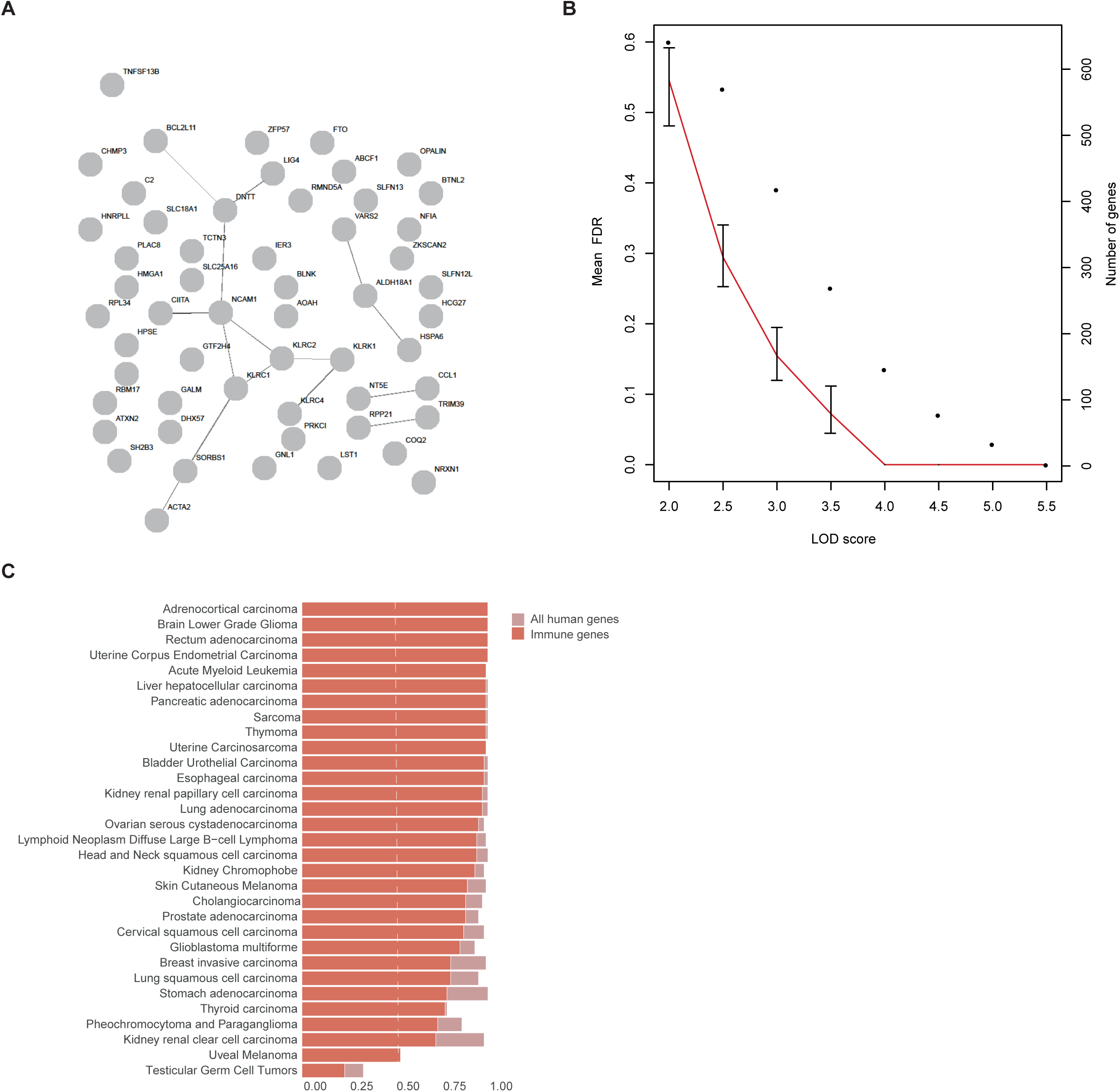
Human GWAS results didn’t form a coherent network, mutational enrichment across genes validated in human GWAS, Related to Figure 5. A. A protein-protein interaction network of identified genetic associations in human GWAS.
B. False discovery rate (red line) for 100 random sets of genes compared, LOD scores lower than the significant threshold are shown.
C. Analysis for 47 CC derived genes validated in human. Percentile is shown for cancer type, the percentile calculated after removing non-immune genes out of genes expressed in immune cell only (red), all human genes (light red).

## Supplementary tables

<u>Table S1</u>- Related to Figure 1: All mouse strains that were included in the study. CC mouse have a typical CC strain identifier; founder strains the full strain name. The table determines at which cohort the mouse was profiled.

<u>Table S2</u>- Related to Figure 1: Detailed list of all CyTOF antibodies (markers) that were used in the study, including clone and vendor. For each antibody indicated whether it was used for clustering purposes, as Citrus input for the calculation of events (cells) distances, and/or mapping, Scaffold uses markers to map clusters to appropriate manual gates based on distance between both.

<u>Table S3</u>- Related to Figure 1: The table indicates for each CC strain in the first cohort of mice which immune cell subsets could be detected in that strain using our phenotypic panel.

<u>Table S4</u>- Related to Figure 1: The table summarize frequency for each population identified in 1. manually gated populations in CC mice bone marrow 2. clustered populations in CC mice bone marrow 3. manually gated populations in human bone marrow.

<u>Table S5</u>- Related to Figure 1: All genes that were tested in QTL mapping process, the genes are expressed in immune cells according to ImmGen and predicted to have a mutation that has moderate or high impact on the protein structure according to Mouse Genome Project available at: http://www.sanger.ac.uk/sanger/Mouse_SnpViewer/rel-1505.

The table also indicates which types of mutations are possible in the gene: single nucleotide polymorphisms (SNPs), insertions/deletions(INDELS), structural variants (SVs).

<u>Table S6</u>- Related to Figure 2: Table of all associations that were detected in the study and met our filtering criteria. For each association (i) left flanking SNP of the genetic interval is shown, (ii) the chromosome on which located the genomic loci (iii) it’s actual position on the chromosome in cM (iv) LOD score of the association (v) phenotype to which the association was made (vi) Gene that is included in the interval (vii) effect size of the gene.

<u>Table S7</u>- Related to Figure 2: Summarize the total number of associations with each cell subset.

<u>Table S8</u>- Related to Figure 2: The table summarize frequency for each population identified in manually gated populations in CC mice bone marrow of second cohort of mice.

<u>Table S9</u>- Related to Figure 2: The table indicates for each CC strain in the second cohort of mice which immune cell subsets could be detected in that strain using our phenotypic panel.

<u>Table S10</u>- Related to Figure 2: The table shows adjusted *p*-*value* of the validation for each trait-gene association in the second cohort of mice.

<u>Table S11</u>- Related to Figure 2: The table shows mouse phenotypes according to Mammalian Phenotype Ontology for part of genes that were found as associated to immune cell subset frequencies in our study.

<u>Table S12</u>- Related to Figure 3: Results of the enrichment analysis for mouse study by IPA. All significant categories after multiple testing correction are shown. The specific category, disease or functional annotation, global annotation (manually curated), the actual genes included, number of genes in the category and the adjusted p-value of the hypergeometric test.

<u>Table S13</u>- Related to Figure 4: Table shows percent of IdU positive cells for each strain in each cell subset.

<u>Table S14</u>- Related to Figure 5: Results of the enrichment analysis for three human GWAS by IPA. All significant categories after multiple testing correction are shown. The specific category, disease or functional annotation, global annotation (manually curated), the actual genes included, number of genes in the category and the adjusted p-value of the hypergeometric test.

<u>Table S15</u>- Related to Figure 5: List of genes found in three human studies: Orrù *et al.*, 2013; Roederer *et al.*, 2015; Patin *et al.*, 2018.

<u>Table S16</u>- Related to Figure 5: Table shows genes that were found in mice and validated in human GWAS. For each gene we indicated association p-value, the immune phenotype it associated with and its name as it appears in the original paper Roederer *et al.*, 2015.

